# Derivation of Genetically-Defined Murine Hepatoblastoma Cell Lines with Angiogenic Potential

**DOI:** 10.1101/2025.07.25.666764

**Authors:** Keyao Chen, Ahmet Toksoz, Colin Henchy, Jessica Knapp, Jie Lu, Sarangarajan Ranganathan, Huabo Wang, Edward V. Prochownik

**Author notes:** Co-Correspondence: EVP; HW.

## Abstract

**Background/Objectives:** Hepatoblastoma (HB) is the most common form of pediatric liver cancer, with the vast majority of these tumors evidence of mutation and/or deregulation of the oncogenic transcription factors β-catenin (B), YAP (Y) and NRF2 (N). HB research has been hampered by a paucity of established cell lines, particularly those bearing these molecular drivers. All combinations of B, Y and N (i.e. BY, BN, YN and BYN) are tumorigenic when over-expressed in murine livers but it has not been possible to establish cell lines from primary tumors. Recently, we found that concurrent Crispr-mediate targeting of the *Cdkn2a* tumor suppressor locus allows for such immor-talized cell lines to be generated with high fidelity.

**Methods:** We generated 5 immortalized cell lines from primary *Cdkn2a*-targeted BN and YN HBs and characterized their properties. Notably, 4 of the 5 retain their ability to grow as subcutaneous or pulmonary tumors in the immune-competent mice from which they originated. Most notably, when maintained under hypoxia conditions for as little as 2 days, BN cells reversibly up-regulated the expression of numerous endothelial cell (EC)-specific genes and ac-quired EC-like properties that benefited tumor growth.

**Conclusions:** The above approach is currently the only means by which HB cell lines with pre-selected, clinically relevant oncogenic drivers can be generated and the only ones that can be studied in immune-competent mice. Its generic nature should allow HB cell lines with other oncogenic drivers to be derived. A collection of such cell lines will be useful for studying tumor cell-EC trans-differentiation, interactions with the immune environment and drug sensitivities.

**Simple Summary:** Most hepatoblastomas (HB) are associated with aberrant expression of β-catenin (B), YAP (Y) and/or NRF2 (N) transcription factors and can be modeled in mice by over-expressing pairwise of triple combination of these. Virtually no human or murine HB cell lines exist that bear these mutations. We describe here an efficient way to generate cell lines from primary BN and YN tumors. Moreover, one of the BN lines shows a remarkable ability to trans-differentiate into endothelial cells under hypoxic conditions that may facilitate angiogenesis. These cell lines along with previousl-derived BN and BYN lines showed similar sensitivities to drugs commonly used to treat HB. Because the approach for cell line derivation we describe is quite general, it should allow for the generation of additional lines driven by less common factors. A collection of such permanent and well-characterized cell lines will facilitate studies that are difficult or impractical to perform *in vivo*.

## 1. Introduction

Hepatoblastoma (HB), the most common pediatric liver cancer, arises almost exclusively in children less than 4 years of age and is classified into several distinct histopathologic subtypes [1,2]. Despite this complexity, it harbors fewer recurrent genetic alterations than any other known childhood or adult malignancy [3,4]. In this regard, >70% of HBs carry missense or in-frame deletion mutations in the *CTNNB1* gene, which encodes the β-catenin transcription factor (TF) and ∼50% carry missense mutations in or amplify the NRF2 gene, which encodes the NRF2 TF [5-12]. These changes dysregulate the cytoplasmic-to-nuclear shuttling of the TFs between the cytoplasm and nucleus and result in their constitutive nuclear confinement and target gene deregulation. Under these circumstances, β-catenin’s regulation by Wnt growth factor signaling is lost and NRF2 loses its normal responsiveness to oxidative and xenobiotic stresses [9-11]. A third TF, YAP, is also commonly deregulated in HB [9,13-15]. Along with its close paralog and co-activator, TAZ, YAP is the terminal TF of the Hippo pathway that regulates organ growth and cellular proliferation in response to signals as diverse as cell-cell contact, thrombin and glucagon [16-18]. Although recurrent mutations have noy been identified in the Hippo pathway in HB, as many as 70% of tumors show strong nuclear localization of YAP [9,13-15]. Finally, up to 50% of human HBs show evidence for hypermethylation and/or down-regulation of the CDKN2A tumor suppressor locus, which encodes the tumor suppressors (TSs) p16^INK4A^ and p19^ARF^ in partially overlapping reading frames [19-26].

Mouse models of HB have demonstrated that patient-derived and nuclear-localized mutations of β-catenin such as the 90 amino acid in-frame deletion Δ90 (hereafter B) and the NRF2 missense mutation L30P (N) as well as a nuclear-localized S127A missense mutation of YAP (Y), can induce HBs when expressed in the liver in any pair-wise combination (BY, NY and BN) and that the triple combination (BYN) generates particularly aggressive tumors [10,12,15,27]. Each of these tumor groups, which can be induced via the hydrodynamic tail vein injection (HDTVI) of Sleeping Beauty (SB) vectors encoding the above-mentioned mutant TFs, demonstrates distinct features. For example, BY tumors are associated with a median survival of ∼90 days and histologically resemble the most common “crowded fetal” HB subtype [10,12,15,27]. BN and YN combinations generate tumors associated with a 2-fold longer median survival with the former being more differentiated and the latter more resembling undifferentiated hepatocellular carcinoma (HCC) with HB-like features [11]. Finally, BYN tumors, with a median survival of <30 days, also display the crowded fetal pattern but possess innumerable and highly characteristic fluid-filled cysts that often abut well-demarcated foci of necrosis [11,26].

Few established human HB cell lines currently exist and even fewer are readily available; moreover, some, such as HepG2, originate from atypical patients and/or possess unusual molecular features that are not representative of the vast majority of human tumors [28,29]. Similarly, murine cell lines established from *MYC* oncogene-driven tu-mors more closely resemble the undifferentiated HCCs with HB-like features originally described in mice bearing tumors driven by a doxycycline-inducible human *MYC* transgene [30-32]. Importantly, these Myc-driven tumors and cell lines contain none of the common HB-associated mutations described above. Additionally, Myc is not necessary for HB initiation since its genetic ablation in hepatocytes still allows for highly efficient B+Y-generated tumorigenesis [10,27]. These findings underscore the need for HB cell lines that are molecularly defined, are driven by clinically relevant oncogene and/or TS combinations and that represent different histopathologic variants. The availability of such cell lines, which could be reliably generated from murine HBs, would make them useful for *in vitro* genetic manipulation and/or studies requiring controlled and/or rapid changes in the extracellular environment. They would also provide ideal reagents with which to screen for new chemotherapeutic drugs and to determine how their susceptibility to these or more traditional agents is impacted, if at all, by the underlying molecular drivers. Finally, these cell lines could potentially be propagated as orthotopic, subcutaneous and/or metastatic tumors in the immuno-competent mice from which they originate, thus permitting studies of host-tumor interactions that are otherwise impossible to conduct with human cell lines [26].

Having failed to generate immortalized cell lines from any of the above-described B-, Y- and/or N-driven HB types in over 30 prior attempts, we recently succeeded in efficiently deriving 8 BY and 3 BYN murine HB cell lines [26]. This was made feasible by using *in vivo* Crispr/Cas9-mediated gene editing to mutate the *Cdkn2a* TS locus at the time of tumor initiation [26]. We have shown that these cell lines as well as primary tumors can be markedly growth-inhibited when wild-type (WT) p16^INK4A^ or p19^ARF^ expression is restored thus attesting to their importance in tumorigenesis [26,33]. Ten of the above 11 cell lines were tumorigenic and retained the appearance of HBs when propagated subcutaneously in the immunocompetent FVB/N mouse strain from which they originated [26]. They also grew as pseudo-metastatic lung and lymphatic tumors when administered to recipient mice via standard tail vein injections, could be maintained *in vitro* as tumor spheroids on non-adherent surfaces and, in doing so, acquired hypoxic interiors, with some reversibly acquiring the properties of endothelial cells (ECs) [26].

We now describe the derivation and characterization of 5 additional immortalized HB cell lines originating from the more slowly growing BN and YN primary tumor types [26]. As with BY and BYN cell lines, the derivation of BN and YN cell lines also required concurrent *Cdkn2a* inactivation, which is often a feature of HBs [19-26]. Four of the cell lines could also be propagated as subcutaneous and metastatic tumors. Like BY and BYN cell lines, BN and BY cells expressed EC markers and assumed EC-like behaviors to variable degrees, either when allowed to form spheroids on non-adherent surfaces or when propagated as monolayer cultures under hypoxic conditions. When combined with BY and BYN cell lines, BN and YN cell lines could also be used to assess their relative sensitivities to 4 of the chemotherapeutic drugs most commonly used to treat HB.

## 2. Materials and Methods

### 2.1. Animal care and husbandry

FVB/N and nu/nu mice (Jackson Labs, Inc., Bar Harbor, ME) mice were housed in micro-isolator cages in a pathogen-free facility at UPMC Children’s Hospital of Pittsburgh and provided *ad libitum* with standard animal chow and water. All care, diets and procedures were performed in accordance with the Public Health Service Policy on Humane Care and Use of Laboratory Animal Research Guide for Care and Use of Laboratory Animals. All experimental details have been previously described and were approved by the University of Pittsburgh’s Institutional Animal Care and Use Committee (IACUC) [26].

### 2.2. Plasmids, plasmid DNA purification, hydrodynamic tail vein injections (HDTVIs) and transfections

Sleeping Beauty (SB) vectors encoding a patient-derived 90 amino acid in-frame deletion (Δ90) of β-catenin (B), a patient-derived missense mutation (L30P) of NRF2 (N) and a missense mutation (S127A) of the Hippo pathway terminal TF YAP (Y), have been previously described and their pair-wise and triple oncogenic potentials demonstrated [11,12,15,27]. HDTVI inocula were administered to 4-6 wk old FVB/N mice over ∼5 sec. in 2 ml of PBS. These contained 10 μg each of B+N or Y+N SB vectors, 2 μg of a non-SB vector encoding SB transpose and 2 μg each of 2 pDG458 Crispr/Cas9 vectors [27] each of which encoded 2 gRNAs (C1+C2 and C3+C4) directed against exon 2 of the murine *Cdkn2a* gene [26]. For *in vitro* transfection of HB cell lines, pSBbi-RP SB vectors encoding dTomato and puromycin N-acetyl transferase (Addgene, Inc., Watertown, MA) along with wild-type (WT) or relevant mutant forms of p16^INK4A^ and p19^ARF^ derived from BY and BYN cell lines (2 μg each, plus 0.5 μg of the SB transposase) [27] were stably introduced into the indicated tumor cell lines using the Superfect transfection reagent according to the directions of the supplier (Thermo Fisher, Inc. Pittsburgh, PA).

To readily monitor the acquisition of endothelial cell (EC)-like properties, some cell lines were transfected *in vitro* with 2 μg of a previously described vector encoding EGFP under the control of the EC-specific Tie2/Tek promoter (Tie2-EGFP) [34]. Stable clones were selected in Geneticin (G418, ThermoFisher) (400 μg/ml) and then pooled for all subsequent studies.

#### 2.3. Establishment of immortalized BN and YN cell lines

Immortalized cell lines were derived as described previously for BY and BYN cell lines [26]. Briefly, upon reaching maximum allowable size, BN and YN tumors (each harboring multiple *Cdkn2a* mutations) were excised, minced into 1-2 mm pieces and washed 3 times in PBS. After digesting for 30 min in 0.1% trypsin at 37°C (Sigma-Aldrich, Inc. St Louis, MO), the tissue fragments were further dispersed by vortexing and vigorous pipetting and were then incubated in 4-5 100 mm tissue culture plates containing Dulbecco’s modified D-MEM+10% FBS that was supplemented with L-glutamine and penicillin G/streptomycin (Corning Life Sciences, Corning, NY). Over the next several days-weeks tumor cells gradually attached to the surface of the plates but did not replicate. Subsequent to this, cells were trypsinized weekly, with those remaining adherent to the plates being discarded. Eventually, small colonies of non-contact-inhibited cells appeared and were expanded further.

### 2.4. Growth curves

2000 viable tumor cells were seeded into individual wells of 12-well plates and maintained under standard growth conditions as previously described [26]. Counts were performed at regular intervals on 3-4 replicas using an Incucyte SX5 live cell imaging instrument (Sartorius Instruments, Göttingen, Germany).

### 2.5. Tumorigenicity of immortalized YN and BN cell lines

YN or BN cell monolayers were trypsinized, washed twice in PBS and re-suspended in PBS at a concentration of 5×10^6^ cells/ml. 0.2 ml were delivered via subcutaneous injection into the flanks of 4-6 wk old FVB/N or nu/nu mice. To determine the “pseudo-metastatic” potential of the cell lines the same number of cells was delivered via slow tail vein injection. Mice were observed twice weekly for the appearance of palpable tumors. Tail vein-injected mice were sacrificed after 6-8 wks or before if noted to be in any distress.

### 2.6. Tumor histology and confocal microscopy

Subcutaneous and lung tumors were excised, immediately fixed in PBS-4% paraformaldehyde and then processed for H&E staining at the UPMC Children’s Hospital of Pittsburgh Core Histology Facility using standard procedures [12,26]. EGFP in tumor sections was examined following the embedding of fresh tumor tissue in O.C.T. compound (Thermo Fisher). After straining with Hoechst 33342, fluorescence imaging was performed with an Olympus Fluoview 1000 confocal microscope (Tokyo, Japan).

### 2.7. Tumor cell responses to hypoxia

The effects of hypoxia on tumor cells were tested in 2 different ways. First, logarithmically growing tumor cells were trypsinized, re-suspended at a concentration of ∼10^6^ cells/ml in DMEM-10% FBS and re-seeded into 15 ml polypropylene tubes (2 ml/tube). The tubes were incubated under normoxic conditions (20% O_2_ + 5% CO_2_) in a standard tissue culture incubator at a 30° angle with loosely attached caps that maintained sterility while allowing for gas exchange. Under these conditions, tumor spheroids formed rapidly and could be maintained in a highly viable state for at least 1-2 weeks. Small numbers of spheroids were removed periodically for staining with the hypoxia sensitive dye Image-IT Green (ThermoFisher) as previously described [26]. The same cells, stably transfected with the above-described Tie-2-EGFP vector [34], were also periodically examined by fluorescence microscopy for the expression of EGFP as a general surrogate for the expression of EC-specific genes. In other studies, the above cells were cultured as monolayers under normoxic conditions for 1-2 days and then exposed to an atmosphere of 1% oxygen + 5% CO_2_ in a Heracell Vios 160i CO_2_ incubator (Thermo Fisher) at 37°C for the indicated periods of time followed by image-IT Green staining or imaging for EGFP expression.

### 2.8. Isolation of EGFP+ cells

Monolayer cultures of BN cells lines stably transfected with Tie2-EGFP in early log-phase growth were exposed to 1% oxygen for 48 hr as described above. Cells expressing the highest levels of EGFP were then isolated by fluorescence-activated cell sorting using a Becton-Dickinson FACSAria III instrument (Franklin Lakes, NJ) and then further expanded under normoxic conditions.

### 2.9. SDS-PAGE and Immunoblotting

Livers, tumors and cells were washed in PBS, snap-frozen in liquid nitrogen and stored at -80°C until being further processed and analyzed as described previously [26].

Standard SDS-lysis buffer was prepared in the presence of protease and phosphatase inhibitors according to the directions of the supplier (ThermoFisher) and as previously described [12,26,27]. Tissues were rapidly homogenized in a Bullett Blender (Stellar Scientific, Baltimore, MD), diluted 1:1 in 2 x SDS-PAGE running buffer, boiled for 10 min and then clarified by centrifugation before storing at -80C in small aliquots. Immuno-blotting was performed using PVDF membranes and semi-dry transfer conditions [26,27]. Antibodies used included those directed against p16^INK4A^ (#Ab211542, 1:2000, Abcam, Cambridge, UK), p19^ARF^ (#NB200-174, 1:1000, Novus Biologicals, Cantennial, CO), GAPDH (#G8795, 1:10000, Sigma-Aldrich), YAP (#4912, 1:1000, Cell Signaling Technologies [CST], Inc., Danvers, MA). Additional antibodies included horseradish-peroxidase (HR)-conjugated goat anti-mouse IgG (#7076, 1:10000, CST), HRP-goat anti-rabbit IgG (#7074, 1:5000, CST), and HRP-goat anti-rat IgG (#7077, 1:5000, CST). All immuno-blots were developed using a Pierce Enhanced ECL Chemiluminescence Detection Kit according to the directions of the vendor (ThermoFisher, Inc., Pittsburgh, PA). Signal relative to GAPDH was quantified by density scanning using ImageJ software.

### 2.10. *Cdkn2a* exon 2 amplicon sequencing

Briefly, DNA was extracted from cell lines harboring *Cdkn2a* mutations using DNeasy columns according to the directions of the supplier (Qiagen, Inc., Germantown, MD). Exon 2 of the *Cdkn2a* locus was amplified by PCR using different barcoded primers and the previously described conditions [26]. PCR products were combined and subjected to deep sequencing using the Illumina MiSeq platform (Azenta Life Sciences, Inc., Chelmsford, MA). Sequencing reads were imported into CLC Genomics Workbench 24 (Qiagen), merged and de-multiplexed into individual groups that were determined by the identities of the bar-coded primers. Individual reads from each group were then aligned to the mouse *Cdkn2a* locus.

### 2.11. RNAseq and bio-informatics studies

Total RNAs were isolated as previously described using RNeasy columns (Qiagen). Only samples with RIN values >8.0 were processed further. Library preparation and sequencing were performed at the Pittsburgh Liver Research Center’s Core Sequencing laboratory (https://livercenter.pitt.edu/gsbc/#Sequencing) using an AVITI/Illumina instrument. Analyses were performed using nf-core/rnaseq version 3.12.0 as previously described [26]. Genes with low expression (average counts among all groups < 1) were filtered out. Normalized counts were generated by DESeq2 and then used for statistical quantification and heat map generation. Gene set enrichment analysis (GSEA) was performed using the clusterProfiler R package. Endothelial gene sets for GSEA analysis were achieved by filtering all “C8” gene sets from Molecular Signatures Database (MSigDB) with “ENDOTHELIAL” in their names. Data from The Cancer Genome Atlas Program (TCGA) were accessed via UCSC Xena browser with batch-normalized TPM used for quantification. Only samples with both survival and RNA-seq data were selected for analyses. Principal Component Analysis (PCA) and k-means clustering were performed to sub-classify tumors into different clusters. Survival curves were generated using the survival R package. P-values were calculated using log-rank test and adjusted with Benjamini-Hochberg (BH) method for multiple comparisons.

### 2.12. Drug sensitivity studies

Monolayer cultures were trypsinized and 3×10^3^ cells, in a volume of 100 μl, were seeded in D-MEM-FBS into individual wells of 96 well plates. The following day fresh medium containing the indicated concentrations of drugs was added for 72 hr at which point standard MTT assays were performed as previously described (6 replicas/dose) [35-37]. Assays were read on a SpectraMAX Plus Micro Plate Reader (Molecular Devices, Inc. San Jose, CA) spectrophotometer. Drugs used included cis-platinum (Santa Cruz Biotechnology Inc.), etoposide (Sigma-Aldrich), doxorubicin (Sigma-Aldrich) and vincristine (Sigma-Aldrich).

### 2.13. Statistical analyses

R software v4.4.0 (R Foundation for Statistical Computing, Vienna, Austria) and GraphPad Prism v9.00 (Dotmatics, Inc. Boston, MA) were used for statistical analyses. The ComplexHeatmap package was utilized for heatmap visualizations. The number of samples per group (n) for each experiment is indicated either in the figure legend or within the figure itself. Two-tailed, unpaired t-tests were used to compare significance between normally distributed populations and two-tailed Mann-Whitney exact tests were used to determine the significance between non-normally distributed populations.

## 3. Results

### 3.1 Efficient generation of BN and YN cell lines

To derive immortalized cell lines from primary BN and YN HBs, we relied upon a previously used strategy that permitted the generation of 11 BY and BYN cell lines with 100% efficiency [26]. For this, Sleeping Beauty (SB) vectors encoding B+N or B+Y were delivered via HDTVI along with a non-SB vector encoding SB transpose [12,26]. While these 2 combinations alone generated tumors with >95% efficiency, key to the establishment of immortalized cell lines was the inclusion of non-SB-based Crispr/Cas9 vectors that allowed for mutagenesis of exon 2 the *Cdkn2a* locus [26]. *Cdkn2a* encodes the p16^INK4A^ and p19^ARF^ TS proteins in overlapping reading frames and is silenced in a significant fraction of human HBs [24-26,38]. Re-expression of either of these TSs in BY and BYN cell lines or primary tumors is also markedly growth-suppressive [26,33]. The gross appearances of the BN and YN tumors generated by this method were indistinguishable from those described previously in the absence of *Cdkn2a* targeting (Figure 1A) [11]. Examination of HJ&E-stained sections showed that all tumor histologies resembled those previously reported, thus indicating that *Cdkn2a* inactivation also did not significantly affect the microscopic appearance of tumors (Figure 1B) [11]. BN tumors also contained a somewhat more extensive vascular network (Figure 1B).

**Figure 1.**
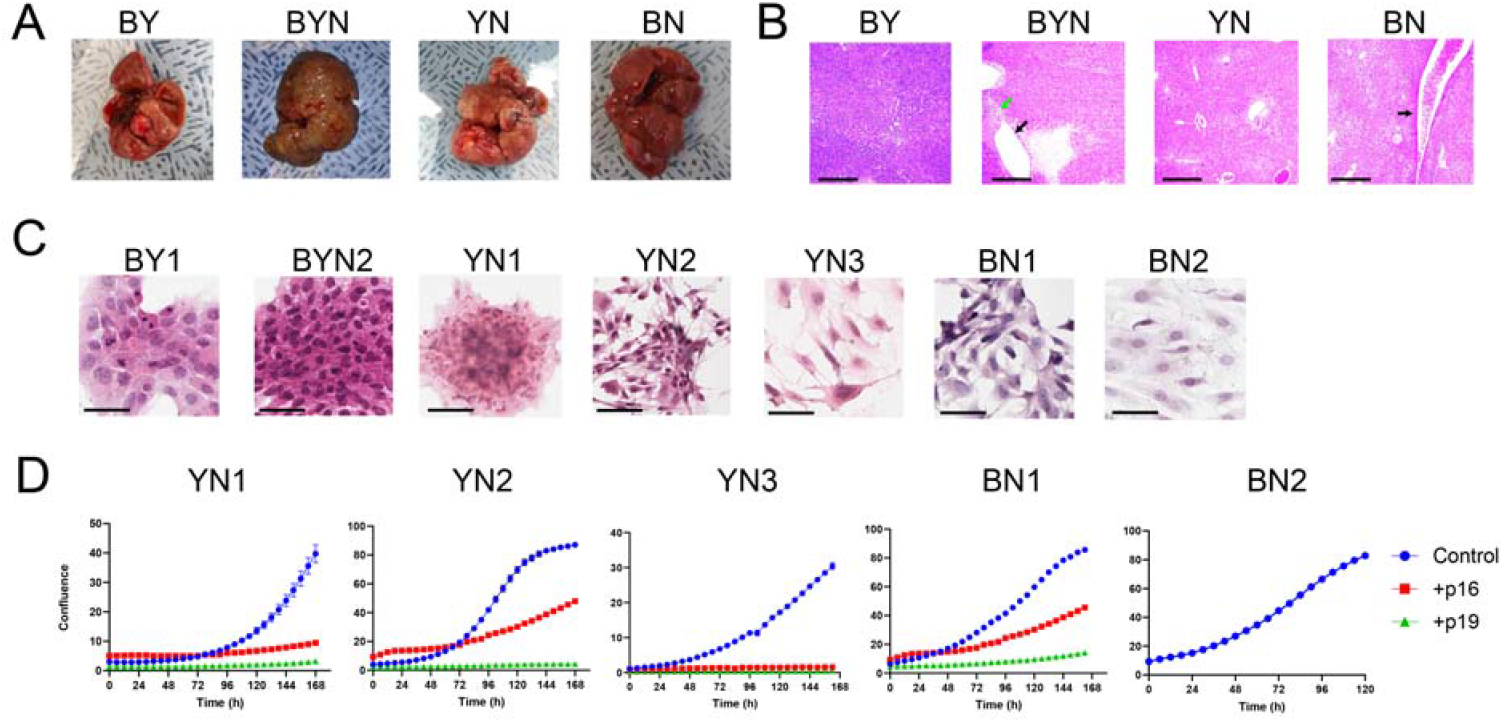
Properties of BN and YN HBs and their progeny immortalized cell lines. **(A)** Gross appearance of typical tumors generated by the indicated combinations of oncogenic drivers. Images of previously generated BY and BYN tumors [26] are included for comparison. To allow for the establishment of immortalized BN and YN cell lines, all tumors were generated with the inclusion of 2 Crispr/Cas9 vectors encoding 4 different gRNAs directed against exon 2 of the *Cdkn2a* locus [26]. **(B)** H&E-stained sections of tumors from **(A)** Note the more prominent blood vessels in BN tumors (black arrow) and the previously reported fluid-filled cysts adjacent to regions of necrosis in BYN tumors (black and green arrows, respectively) [11,26]. Scale bars = 500 μm. **(C)** H&E-stained cells from the indicated immortalized cell lines propagated on coverslips *in vitro*. BY1 and BYN2 cell lines were derived and characterized previously and are included here for compararive purposes [1]. Scale bars = 50 μm. **(D)** Growth curves of the indicated cell lines and their suppression via the enforced expression of WT p16INK4A and p19ARF. The indicated cell lines were transfected with a control pSBbi-RP SB vector or with vectors encoding WT p16INK4A or p19ARF. Two days later, the cells were seeded into 12 well plates and maintained in 2 μg/ml of puromycin while monitoring dTomato expression. Subsequent growth was monitored using an Incucyte S3 Live-Cell imaging and Analysis System. Each point represents the mean of 4 replicas +/-1 S.E. Note that BN2 cells were resistant to transfection on multiple occasions.

As reported for BY and BYN tumors harboring *Cdkn2a* mutations [26], 2 BN and 3 YN cell lines were readily established after 4-8 wks of *in vitro* maintenance. Culturing these on coverslips *in vitro* showed that, with the exception of YN1 cells, which resembled previously described BY and BYN cells, BN and YN cells had a more epithelial appearance (Figure 1C) [1]. All 5 cell lines displayed similar doubling times (25-30 hr.) and the growth of 4 could be markedly suppressed by enforcing the expression of wild-type (WT) p16^INK4A^ or WT p19^ARF^ with the latter being somewhat more potent (Figure 1D). The BN2 cell line could not be examined in this manner as it was resistant to transfection.

### 3.2 BN and YN cell lines express recurrent truncated and/or fused p16^INK4A^ and p19^ARF^ variants that are distinct from those expressed by BY and BYN cell lines

Our previous deep sequencing of BY and BYN HBs revealed a large number of unique p16^INK4A^ and/or p19^ARF^ mutations that were initially generated by *in vivo* Crispr/Cas9 targeting of the *Cdkn2a* locus [26]. However, only a subset of these persisted in tumors and cell lines and an even smaller subset was expressed, mostly in the form of p16^INK4A^ and p19^ARF^ truncations and/or fusions of varying complexities. Similar trends were observed in each of the 5 BN and YN cell lines where we detected an average of 33 unique mutations (range = 29-38), with the top 5 comprising 38-79% of the entire mutational burden (Figure 2A and Supplementary File 1). As with BY and BYN cell lines, immuno-blotting showed unique expression patterns for p16^INK4A^ and p19^ARF^ mutants in each cell line, with only a small number of all possible mutant proteins actually being detected. These ranged from only a single truncated form of p16^INK4A^ in YN1 cells in the absence of any p19^ARF^ to mutant or truncated forms of p19^ARF^ in BN1 cells in the absence of any p16^INK4A^ (Figure 2B).

**Figure 2.**
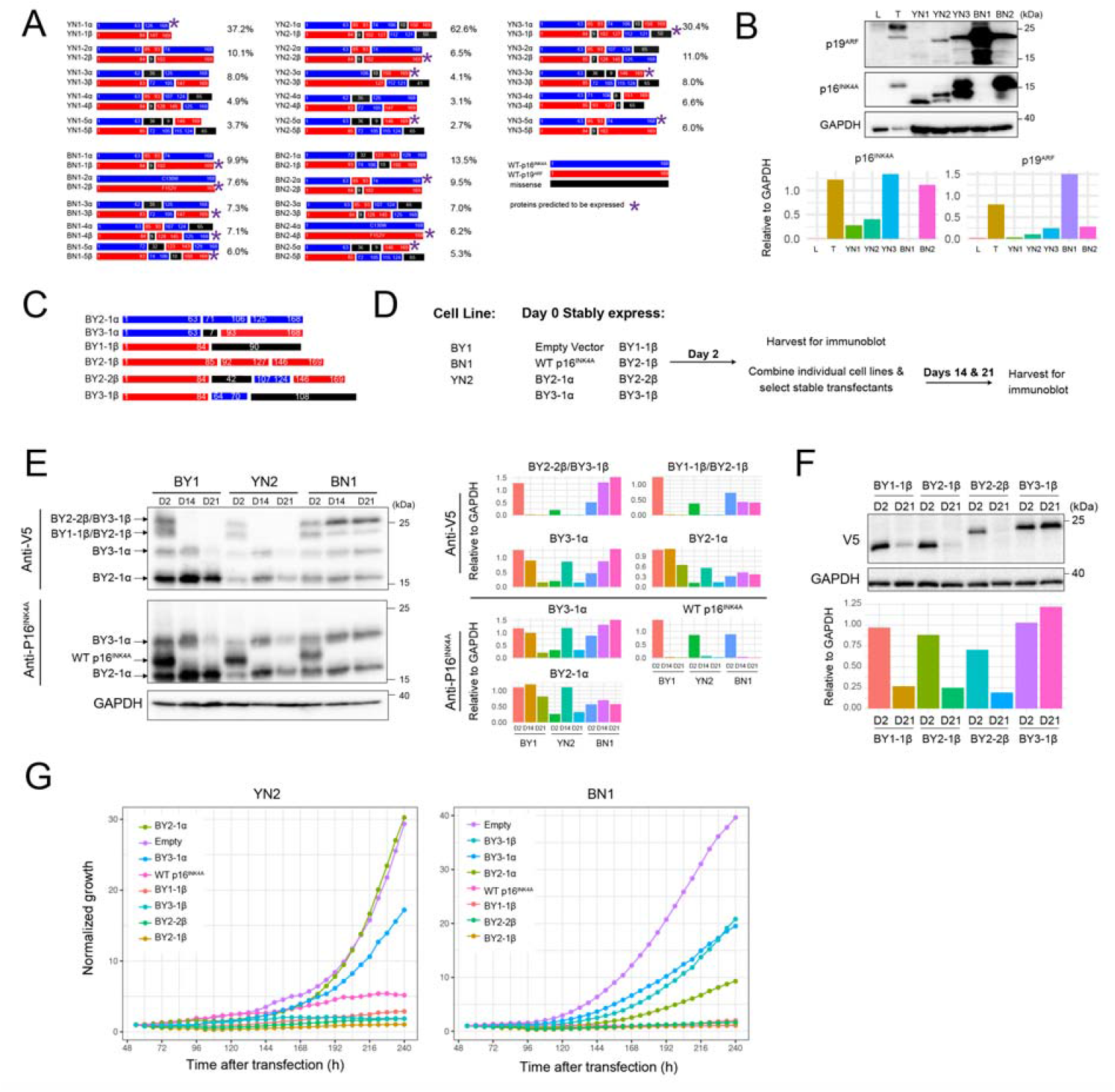
YN and BN cell lines express small, unique subsets of p16^INK4A^ and p19^ARF^ mutants. **(A)** Depictions of the proteins encoded by the 5 most abundant *Cdkn2a* mutations identified in each of the indicated cell lines. The frequencies with which they were detected based on deep sequencing of *Cdkn2a* exon 2 PCR products obtained from cell lines are indicated to the right of each cartoon. Asterisks indicate the mutants that were deemed the most likely to be expressed as proteins, based upon in silico translation of the open reading frames and the sizes of the actual proteins observed by immunoblotting (panel B). Supplementary File 1 contains a more comprehensive list of the mutations identified and their abundance. **(B)** Immunoblots of mutant p16^INK4A^ and p19^ARF^ proteins expressed by the indicated cell lines. Included as controls were a sample of normal liver (L) and a primary BY HB (T) with intact *Cdkn2a* loci. The latter expressed WT p16^INK4A^ and p19^ARF^ as previously described [26]. Expression of each protein relative to that of GAPDH, as determined by densitometric scanning, is shown beneath the blot. **(C)** BY-derived mutant and/or fusion p16^INK4A^ and p19^ARF^ proteins used for expression in BN and YN cell lines. All encoded proteins were V5 epitope-tagged to allow expression levels to be directly compared. See ref. 29 for previous characterization. **(D)** Approach to evaluating the growth suppressive effects of the mutants depicted in C on BN1 and YN2 cells. (E) Differential selection of p16^INK4A^/p19^ARF^ mutants. The indicated cell lines were individually co-transfected with pSBbi SB vectors encoding the mutant proteins depicted in panel **(C)** plus an equal amount of the empty pSBbi vector. Two days later, half the cells were used to assess the transient expression of each protein as described in **(D)**. The remaining cells were selected in puromycin for 14 and 21 days and assessed for the expression of their respective protein at these times using anti-V5 or anti-p16^INK4A^ antibodies. Expression of each protein relative to that of GAPDH, as determined by densitometric scanning, is shown to the right of the blot. **(F)** Selective retention of the non-resolvable mutants shown in E. BN1 cells were separately co-transfected with the empty pSBbi vector alone plus one encoding each of 4 indicated mutants. Puromycin-resistant clones were then selected and expanded for 3 wks as in **(E)** followed by immunoblotting to detect each of the V5-tagged mutants. Expression of each protein relative to that of GAPDH, as determined by densitometric scanning, is shown beneath the blot. **(G)**. The indicated vectors were transfected into YN2 or BN1 cells, which were then seeded into 12-well plates 2 days later in the presence of puromycin and enumerated over the course of the next 10 days. Each point represents the mean of 4 replicas +/-1 S.E.

More so than in BY and BYN cell lines, the *Cdkn2a* derived mutant proteins expressed by BN and YN cells were often recurrent (Figure 2A) [26]. Such examples included the C130W and F152V point mutations seen in both the BN1 and BN2 cell lines as well as the more complex BN2-5β, YN2-5β and YN3-3β mutations. Additionally, none of the mutations selected in BN and YN cell lines matched those previously generated in BY and BYN cell lines. This may have been a result of the reduced complexity of the Crispr/Cas9 vector mix used to target the *Cdkn2a* locus in the former tumors [26] and/or to the oncogenic backgrounds that were differentially permissive for the selection of mutant subsets.

To examine the latter possibility, we relied upon 6 previously described p16^INK4A^/p19^ARF^ mutants that were expressed at high levels in BY or BYN cell lines despite exerting varying degrees of growth suppression when over-expressed (Figure 2C) [26]. Comprised of p16^INK4A^ and p19^ARF^ truncations, fusions and internal deletions, all were V5 epitope-tagged to allow their expression levels to be directly compared (Figure 2C) (26). We transfected these individually into separate cultures of BN1 or YN2 cells and assessed their transient expression 2 days later in half the population with the remaining half being selected in puromycin for 2 or 3 wks (Figure 2D). A separate transfection of BY1 cells served as controls for differential selection of each mutant [26]. In addition, a vector encoding WT p16^INK4A^ was used as a control for negative selection of the vectors whereas an empty vector was used as a neutral selection control and was co-transfected with each of other vectors (Figure 2D) [26,33]. Immunoblotting with antibodies against p16^INK4A^ and the V5 epitope allowed all proteins to be identified 2 days after transfection. As expected, WT p16^INK4A^ was initially expressed at similar levels in all 3 cell lines but was lost in the stably transfected populations upon further passage whereas expression of each of the mutants was lost or retained in cell line-specific ways (Figure 2E). For example, the BY2-1α mutant was retained in all 3 cell lines whereas the BY3-1α mutant was eventually lost in BY1 and YN2 cells but retained in BN1 cells.

The mutants BY2-2β and BY3-1β as well as BY1-1β and BY2-1β encoded proteins of similar sizes that could not be resolved. To identify which, if any, of these were selectively lost or retained in BN1 cells, we performed individual transfections with the empty pSBbi vector and one encoding each of the above 4 fusions proteins and selected these in puromycin for 21 days before repeating anti-V5 immunoblots. These results of this study showed that BY3-1β and BY1-1β were selectively retained whereas BY2-1β and BY2-2β were lost (Figure 2F). Finally, in separate studies, we transfected YN2 and BN1 cells with each of the above vectors and monitored subsequent proliferation in the presence of puromycin selection over the next 10 days. The differential growth suppression by these vectors closely agreed with the previous immuno-blotting experiments (Figure 2G). Collectively, these results emphasize that each cell line displayed a unique response to the suppressive effects of different p16^INK4A^/p19^ARF^ mutants and likely explains why different mutations among the cell lines were selected for in the first place.

### 3.3. Most YN and BN cell lines retain tumorigenic potential

To determine whether BN and YN cell lines retain their oncogenic behaviors following *in vitro* establishment, we injected them subcutaneously (subQ) into the same FVB strain of immuno-competent mice from which they originated. Similar to what we observed for 10 of the original 11 BY and BYN tumor cells [26], all 3 YN cell lines and one BN cell line were also tumorigenic. Histologic examination of these indicated that they retained the appearance of the originally described primary tumors, with BN1 tumors resembling differentiated HBs and YN1-3 tumors having both HB and variable degrees of HCC-like features (Figure 1B and ref. [2]). Unlike primary BN tumors, BN cell subQ tumors were not associated with a more prominent vasculature (Figure 3A and Supplementary Figure 1). When the tumor cells were delivered to the lungs via slow tail vein injection, tumor nodules, with histologies resembling those of the subQ and primary hepatic tumors could be obtained for 2 of the YN cell lines (Figure 3B and Supplementary Figure 2). Thus, YN and BN tumors retain the histologies of the original HBs from which they originate regardless of whether they are propagated as subcutaneous or “pseu-do-metastatic” tumors.

**Figure 3.**
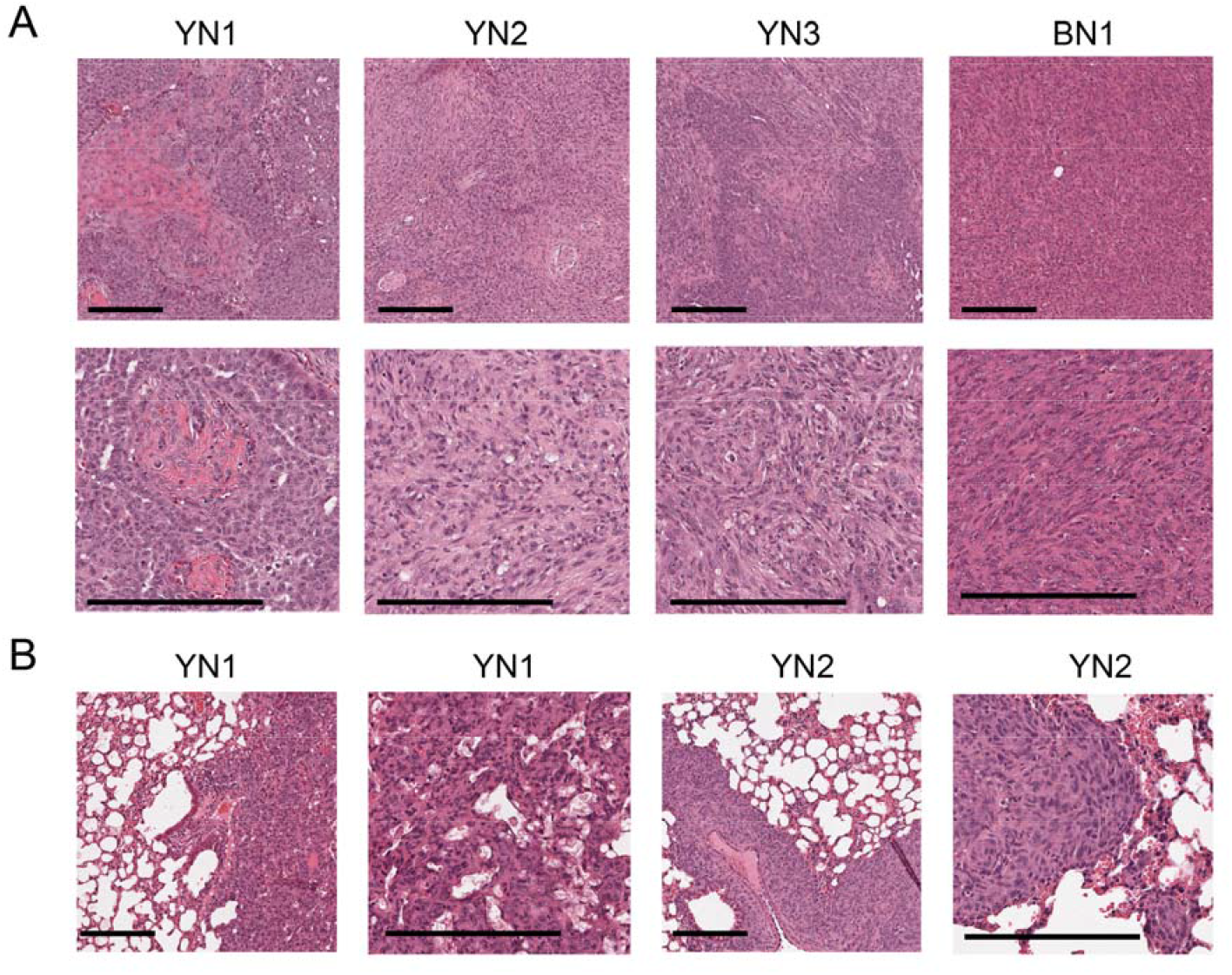
Histologies of BN and YN tumor cell lines following subQ or “pseudo-metastatic” growth in lungs. **(A)** H&E-stained sections of subQ tumors originating from the indicated cell lines. See Supplementary Figure 1 for additional examples. Scale bar: upper panel = 100 μm, lower panel = 200 μm. **(B)** H&E-stained sections of lung tumors generated by tail vein injection of the indicated cell lines. See Supplementary Figure 2 for additional examples. Scale bar: shorter = 100 μm, longer = 200 μm.

### 3.4 BN and YN cell lines form spheroids and acquire endothelial cell-like properties in response to hypoxia

BY and BYN cells were previously shown to form anchorage-independent tumor spheroids that, in response to internal hypoxia, variably up-regulated the expression of an EGFP reporter under the control of the endothelial cell (EC)-specific Tie2/Tek promoter [26,34]. We suggested that this might reflect attempts to re-establish normoxic interiors via the direct “trans-differentiation” of tumor cells into those expressing EC-like properties and potentially capable of generating a neovasculature [34,39-42]. Such EGFP+ cells were previously observed in subQ tumors formed by BY cells and were particularly prominent adjacent to the luminal endothelium of blood vessels [26].

To examine this in BN and YN cell lines, and to allow direct comparisons with our previous results, we first showed that all monolayer cell lines, regardless of their origin, responded similarly to staining with the hypoxia-detecting dye IT-Green after exposure to a 1% O_2_ environment (Figure 4A). However, when the same cell lines were allowed to form spheroids by culturing them on a non-adherent surface for 5 days under normoxic conditions, however, different degrees of IT-Green staining were observed, with the brightest spheroids being those formed by BYN and YN cells (Figure 4B). This indicated that, despite having similar appearances and sizes, the interiors of spheroids from each group likely varied considerably in their degree of hypoxia.

**Figure 4.**
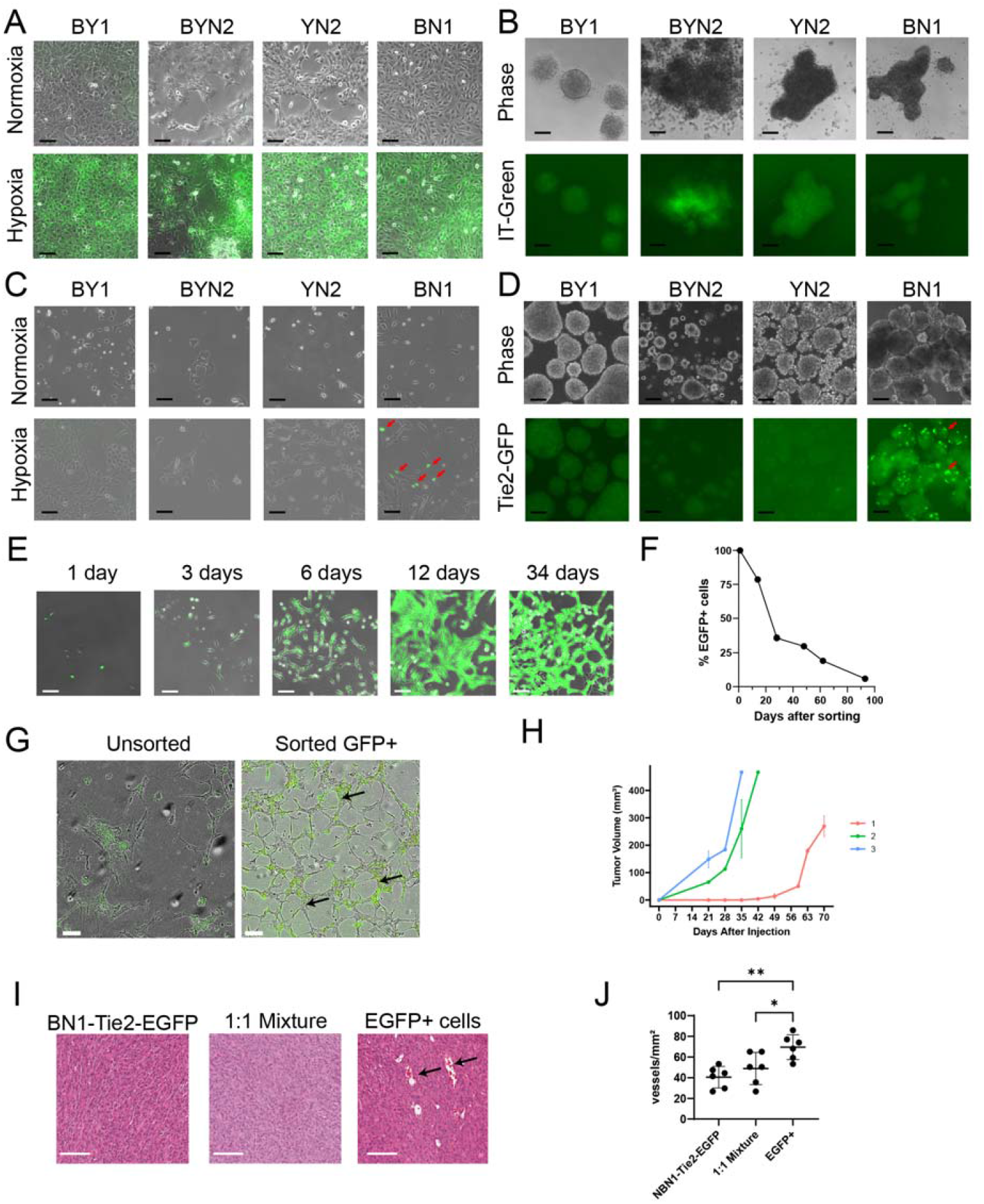
HB cell lines are variably susceptible to EC-like differentiation in response to hypoxia. **(A)** IT-Green staining of monolayer cultures of the indicated cell lines maintained under normoxic or hypoxic (1% O_2_ for 48 hr) conditions. Scale bars = 100 μm. **(B)** IT-Green staining of tumor cell spheroids of the indicated types 5 days after formation. Scale bars = 100 μm. **(C)** Tie2-EGFP expression in the indicated monolayer cultures maintained under normoxic or hypoxic conditions for 48 hr. Arrows indicate EGFP+ cells that were particularly prominent among BN1 cells. Scale bars = 100 μm. **(D)** EGFP expression in the indicated stably transfected Tie2-EGFP cell lines after formation of spheroids for 5 days. Scale bars = 100 μm. **(E)** BN1-Tie2-EGFP cell monolayers were maintained in 1% oxygen for 2 days, at which point, EGFP+ cells were purified by FACS and then expanded and maintained under normoxic conditions for the indicated periods of time. Panels show merged phase contrast and fluorescence images. Scale bars = 100 μm. **(F)** After isolating pure EGFP+ cell populations from hypoxic BN1-Tie2 EGFP cells, they were maintained under normoxic conditions (A). At the times indicated, they were subjected to flow cytometry (ca. 10,000 cells/run) to quantify the percent of EGFP+ cells remaining. **(G)** BN1-Tie2-EGFP cells form EC-like tubes under hypoxic conditions. Left panel: Unsorted BN1-Tie2-EGFP cells were maintained continuously under hypoxic conditions for 6 days. Right panel: Sorted EGFP+ cells from **(E)** were expanded for ∼2 wks under normoxic conditions, re-plated and then maintained under hypoxic conditions for 6 days. Both panels show merged phase contrast and fluorescence images. Scale bars = 100 μm. Arrows indicate representative complete tubes that were formed by the latter group of cells. **(H)**. BN1-Tie2-EGFP, maintained under normoxic conditions were used to generate control subQ tumors in nu/nu mice. A second group of mice was injected with a 1:1 of the same cells plus an EGFP+ population that was generated following exposure to 1% O_2_ for 2 days. Finally, a third group of tumors was generated with a pure population of EGFP+ BN1-Tie2-EGFP cells. Tumor volumes were quantified weekly. In all 3 cases, a total of 10^6^ cells was injected. **(I)**. Representative H&E-stained sections from the groups shown in **(H)**,Scale bars = 100 μm. (J). Quantification of blood vessel number from sections shown in **(H**). 5-6 separate high power fields from each of the tumor groups was randomly chosen and tumor blood vessels in each was enumerated. **(J)**. Quantification of blood vessel number from the tumors depicted in H and I. 5-6 high-power fields from each group were randomly selected and the blood vessel content of each was enumerated.

None of the above cell lines expressed Tie2-EGFP when maintained as monolayer cultures under normoxic conditions (Figure 4C). Hypoxia, however, induced strong EGFP expression in a subpopulation of BN cells but only faint, if any, EGFP expression in the other 3 cell types. The interiors of BN normoxia-maintained spheroids also showed strong EGFP expression whereas those formed by the other 3 cell lines were again dimmer (Figure 4D). Collectively, these studies indicated that BN cells, more so than lines derived from the other 3 tumor types, were particularly prone to the induction of Tie2-EGFP.

To determine how long monolayer cultures of BN1-Tie2-EGFP cells retained EGFP expression, EGFP+ cells were isolated by fluorescence-activated cell sorting (FACS) following a 2-day exposure to 1% oxygen, expanded under normoxic conditions and evaluated periodically by fluorescence microscopy and flow cytometry. EGFP expression was retained for an extended period of time, with ∼40% of cells remaining positive after one month and ∼20% remaining positive after ∼2 months (Figure 4E and F). EGFP+ cells isolated and expanded for ∼2 wks also formed EC-like “tubes” under hypoxic conditions, even in the absence of a basement membrane matrix that is typically needed to support the generation of these structures (Figure 4G) [34,43,44]. In contrast, unsorted BN1 cells did not form tubes under hypoxic conditions, even though a small number EGFP+ cells again appeared. These observations, coupled with the fact that a relatively small fraction of Tie2-EGFP-transfected BN1 cells could be induced to express EGFP (Figure 4C), strongly suggested that only a small sub-population of tumor cells was capable of acquiring EC-like traits in response to hypoxic stress.

To determine whether any of the above-described Tie2-EGFP-transfected cell lines retained tumorigenic behaviors and could express EGFP *in vivo* as we previously showed for BY1 cells [26], we generated subQ tumors and asked 4 questions regarding the biological behaviors of the EC-like BN1-Tie2-EGFP+ cells. First, could they affect the growth rates of subQ tumors if they were combined with BN1-Tie2-EGFP cells that had not been previously exposed to hypoxic conditions? Second, did a pure population of BN1-Tie2-EGFP+ cells remain tumorigenic? Third, did the presence of larger numbers of EGFP+ cells affect tumor histology and finally, how did the presence of large numbers of EGFP+ cells with EC-like properties affect tumor vasculature? To address these questions we compared the properties of 3 groups of subQ tumors. The first group (control) was generated by injecting a pure population of BN1-Tie2-EGFP cells. The second group was comprised of a 1:1 mix of BN1-Tie2-EGFP cells that had been maintained under normoxic or hypoxic conditions, with the latter group being mostly EGFP+. The third group was comprised exclusively of EGFP+ cells. In answer to our first question, each of the latter 2 groups of HBs grew significantly faster than the control group (Figure 4H). This finding also provided an answer for our second question, which showed that the third group of cell retained marked tumorigenic behavior despite their EC-like appearance. Although histologic examination of these 3 groups showed the tumor cells to be indistinguishable from one another (Figure I), the latter group contained a larger number of blood vessels, many of which were much larger than any of those in the control group (Figure 4I and J). Interestingly, examination of frozen sections from these tumors, again failed to show any EGFP+ cells (not shown). Thus, the simplest interpretation of our findings is that EGFP+ EC-like cells contribute to early blood vessel formation *in vivo* but that EGFP expression and perhaps even other EC-like behaviors are transient and eventually disappear as they do *in vitro* (Figure 4 F). Nonetheless, the presence of a pre-existing EC-like population provides significant growth and vasculogenic benefit to subQ tumor growth.

### 3.5.BN tumors and cell lines activate unique populations of EC-specific transcripts

Tie2-EGFP activation and *in vitro* tube formation by BN1 cells in response to hypoxia (Figure 4E-G) indicated that they likely express EC-specific genes other than Tie2/Tek. We thus examined the enrichment in these cells of 15 EC-specific gene retrieved from a variety of sources and found significant changes of each in at least one of the 4 murine tumor groups (Figure 5A). Moreover, the expression patterns of these gene sets relative to those of livers permitted each tumor group to be distinguished from the others. Similar results were found when only the 1853 non-redundant transcripts from the 3750 members of these gene sets were examined. (Figure 5B, Supplementary File 2). A unique ∼600 member subset of these genes, presumably originating from the liver sinusoidal ECs (LSECs) that normally comprise 15-20% of the organ’s cellular mass (Supplementary File 3) [47,48], was largely down-regulated in 3 of the 4 tumor groups, only to be replaced by numerous other non-LSEC-related sets of EC-specific transcripts from various members of the above-mentioned 15 gene sets. Interestingly, many LSEC-related genes continued to be expressed in BN tumors. These 1853 EC-specific genes were also over-expressed to variable degrees in subsets of 49 human HBs from 2 separate previously published studies (Figure 5C & D) [45,46]. Collectively, these findings indicate that, depending on their underlying genetic drivers, both murine and human HBs co-opt EC-specific genes from a variety of largely non-LSEC sources while also downregulating LSEC-related transcripts.

**Figure 5.**
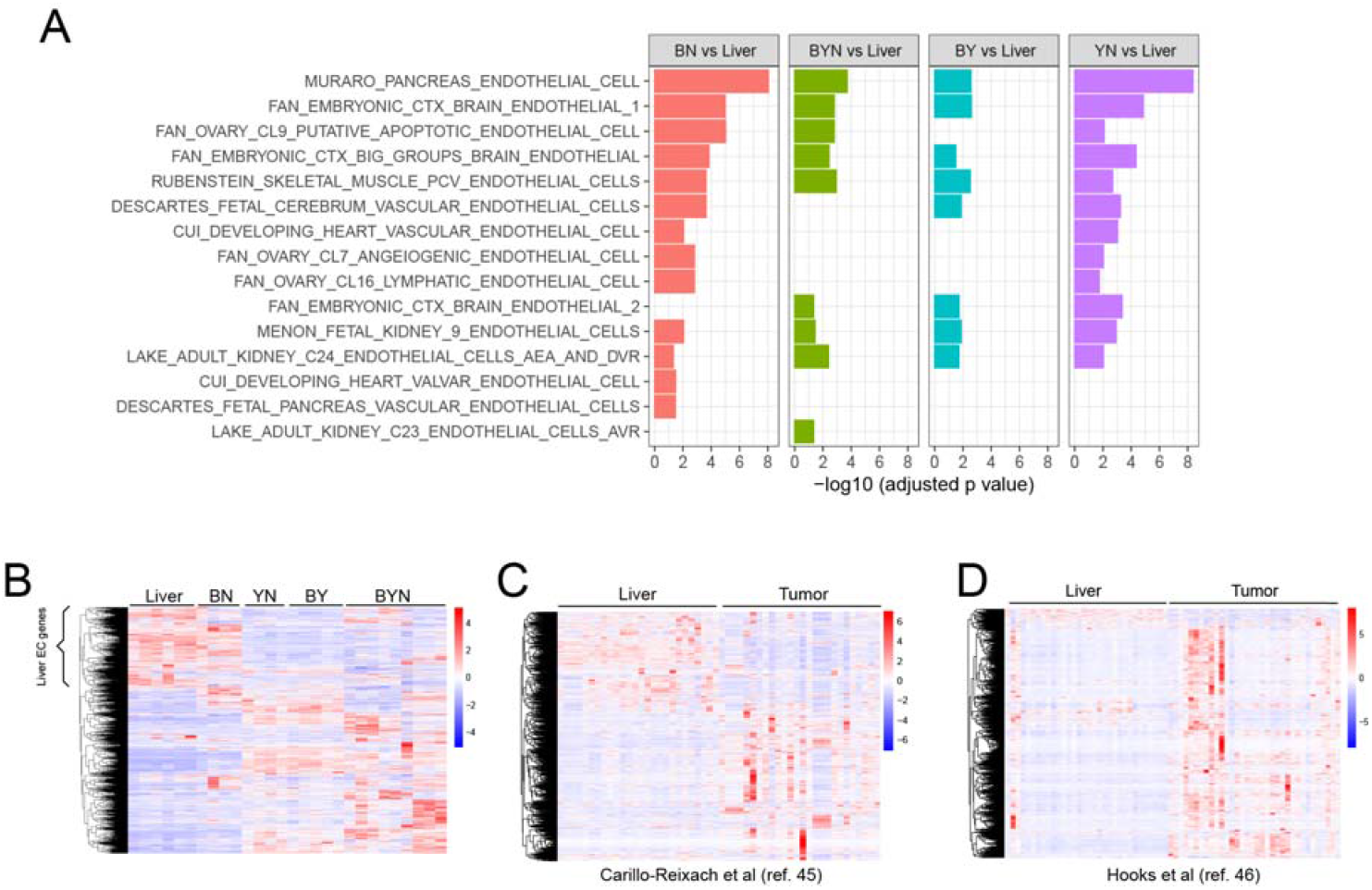
Murine and human HBs up-regulate the expression of EC-specific transcripts. **(A)** Summary of GSEAs performed in primary murine HBs with 15 EC-specific gene collections from the mSigDB containing a total of 3750 transcripts. RNAseq results from our previously reported murine livers and HBs [11,12,27] were used to compare gene expression levels between each of the 4 possible tumor types relative to control normal livers. **(B)** Expression of 1853 unique transcripts extracted from the above 15 EC-specific gene sets in the normal murine livers and tumor groups from **(A)**. The bracket indicates a subset of these transcripts that are likely expressed by LSECs. (C, D) Expression patterns of the human orthologs of the murine genes indicated by the brackets in A genes in 2 independent series of RNAseq studies performed on human HBs [45,46].

To examine the acquisition of EC-like gene expression profiles by tumor cells under more controlled conditions, we exposed unsorted, normoxic monolayer cultures of BN1-Tie2-EGFP cells (“UN”) to hypoxia (1% O_2_) for 2 days (“UH”), expanded the FACS-sorted EGFP+ population for ∼10 days under normoxic conditions (“SN”) and then re-exposed the EGFP+ cells to 1% oxygen for 2 additional days (“SH”) (Figure 6A). Global gene expression profiling revealed 3646-5771 significant differences among the groups (Figure 6B), with many of these being due to anticipated changes in hypoxia-response genes in the UH and SH groups (Figure 6C). Endogenous Tie2/Tek up-regulation, while not appreciated in UH cells due to the relative paucity of those with EC-like properties (Figure 4C) was clearly evident in the sorted populations of SN and SH cells, thus supporting our previous use of EGFP induction as a surrogate marker for EC-like differentiation (Figure 6D).

**Figure 6.**
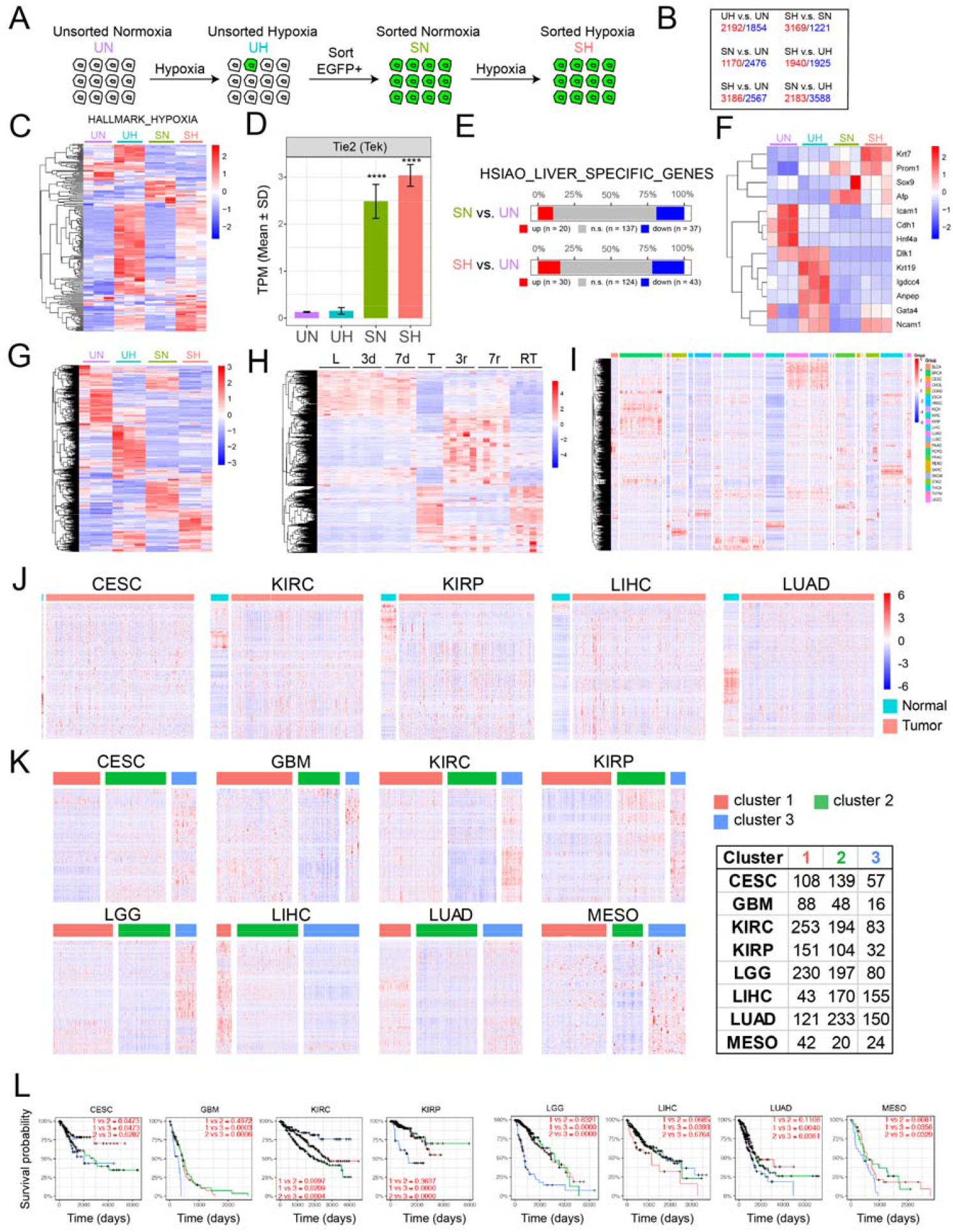
Gene expression profiling of BN1-Tie2-EGFP+ cells reveals widespread EC-specific gene expression within a more undifferentiated sub-population of hepatocytes. **(A)** Scheme for RNAseq profiling of BN1 cells maintained as monolayers under 4 sets of conditions: 1: unsorted cells maintained under normoxia (UN); 2: unsorted cells exposed to hypoxia (1% O_2_) for 2 days (UH); 3: FACS-sorted, EGFP+ cells expanded for ∼14 days under normoxic conditions following a 2 day exposure to hypoxia (SN); 4: SN cells re-exposed to hypoxia for 2 additional days (SH). **(B)** Gene expression differences among BY1-Tie2-EGFP cells cultured under the 4 conditions described in **(A)**. Quantification was performed on 3 replicas of cells from each of the 4 groups. Only genes showing q values <0.05 and fold-differences >2 were included in the analyses. Red numbers indicate genes whose expression was higher in the groups shown at the left; blue numbers indicate genes whose expression was lower in the group shown at the left. **(C)** Heat map of gene expression differences among a group of 200 hypoxia-regulated genes obtained from the mSigDB (Supplementary File 4). **(D)** Expression of Tie2/Tek in each of the 4 cell groups. **(E)** Expression of a group of 194-hepatocyte-specific genes among the indicated groups of cells. **(F)** Expression of 13 genes that are markers of immature hepatocytes and/or hepatocyte precursors [49]. **(G)** Selective expression among the indicated BN1-Tie2-EGFP cell lines of the 1853 EC-specific genes derived from Figure 5B. **(H)** Differential expression of 1853 EC-specific genes from Figure 5B during the course of HCC generation in response to human *MYC* transgene induction [30,32]. L = control liver prior to *MYC* induction; 3d and 7d = days after *MYC* induction and prior to the appearance of any visible tumors; T = HCC after 30 d of *MYC* induction; 3r and 7r = 3 d and 7 d after *MYC* inactivation and the initiation of HCC regression; RT = recurrent HCCs induced ∼3-4 months after the initial tumors had regressed. **(I)** Expression of human orthologs of the 1853 unique EC specific genes from Figure 5 in 23 normal tissues from TCGA. **(J)** EC-specific gene expression from selected normal tissues depicted in (I) along with tumors arising in adjacent regions. **(K)** EC-specific gene expression varies among different subsets of certain human cancers. Principal Component Analysis (PCA) followed by k-means clustering was used to sub-classify each of the indicated types of cancers based on differential levels of expression of the above 1853 EC-specific genes. The list to the right of the heat maps indicates the number of tumors included in each of the indicated categories. **(L)** Kaplan–Meier curves showing the survival of patients among different subgroups in K. Adjusted p-values are displayed for pairwise comparison between different subgroups with Benjamini-Hochberg correction.

Consistent with the idea that SN and SH cells originated from hepatocytes rather than an obscure population of ECs was the fact that they still expressed nearly 200 hepatocyte-specific genes, 10-15% of which were up-regulated (Figure 6E). Notably, these included those encoding α-fetoprotein, Prom1 and other proteins associated with more stem cell-or fetal-like hepatocyte precursors (Figure 6F) [49]. Finally, different subsets of the 1853 EC-specific genes described in Figure 5A were expressed among the 4 different sets of BN1 cells (Figure 6G).

To determine whether the dysregulation of the above 1853 EC-specific genes occurred in other tumors, we first examined their expression in highly undifferentiated murine HCCs that were generated in a rapid and reversible manner by the doxycy-cline-mediated induction of a human *MYC* transgene [30,32]. As observed previously in HBs (Figure 5B), ∼600 of these transcripts were expressed in control livers and did not change significantly for the first 7 days after *MYC* induction, which was well before the appearance of tumors (Figure 6H). Large tumors that were sampled ∼30 days after *MYC* induction showed a dramatic down-regulation of these transcripts and the up-regulation of a new set of previously non-expressed EC-specific transcripts. These largely disappeared 3 days and 7 days after*MYC* silencing, a time of marked tumor regression and tissue re-modeling [30,32] and were replaced by a new set of ectopic EC-specific transcripts. The re-induction of HCCs 3-4 months after the initial tumors had completely regressed was associated with a pattern of EC-specific gene expression closely resembling that of the original tumors. From these studies we conclude that 3 distinct patterns of EC-specific transcript expression are associated with the development and regression of *MYC-*driven HCCs. The first is associated with the previously-described LSEC-specific transcripts of normal livers [45,46]. The second is associated with actively growing tumors and largely replaces the previous transcripts group. Finally, there exists an additional unique set that is expressed only transiently during tumor regression.

We next examined the expression of the above EC-specific transcripts in various normal human tissues and matched tumors from The Cancer Genome Atlas. In the former samples, we determined that, as in the case of murine livers (Figure 5), specific subsets of the above EC-specific genes were expressed in highly tissue-specific ways as expected from previous reports [47,48,50-52] (Figure 6I). In those cases for which a sufficient number of normal tissues could be evaluated, these expression patterns also differed significantly from those of tumors arising in adjacent regions just as they had in murine HBs and HCCs (Figure 6J). Further refinement of these data from 8 different tumor types showed that all could also be further subdivided into those with high, intermediate and low levels expression of their specific transcripts (Figure 6K). In 6 of 8 cases, the subsets with the highest levels of EC-specific gene expression were associated with significantly shorter long-term survival (Figure 6L).

### 3.6. Y suppresses EC differentiation

Speculating that BN cells more readily acquire EC-like properties because Y suppresses EC differentiation [53,54], we co-transfected freshly-sorted BN1-Tie2-EGFP+ cells (Figure 4E) with an empty (control) pSBbi SB vector or one expressing Y. Among the stable transfectants from the latter group, we found many fewer examples of EGFP + Y co-expressing cells as determined by fluorescence microscopy (Figure 7A). Quantification of the dTomato-positive cells by flow cytometry showed a 20-fold reduced intensity of EGFP co-expression (Figure 7B). Additionally, when dTomato^high^ cells from this experiment were sorted, expanded and examined for Y expression by immuno-blotting, they were found to express higher levels of exogenous Y than the unsorted population (Figure 7C). Consistent with previous reports that YAP negatively regulates both itself and the paralogous TF TAZ [55,56], endogenous Y levels were reduced in both populations of cells that expressed Y relative to those transfected with the control vector.

**Figure 7.**
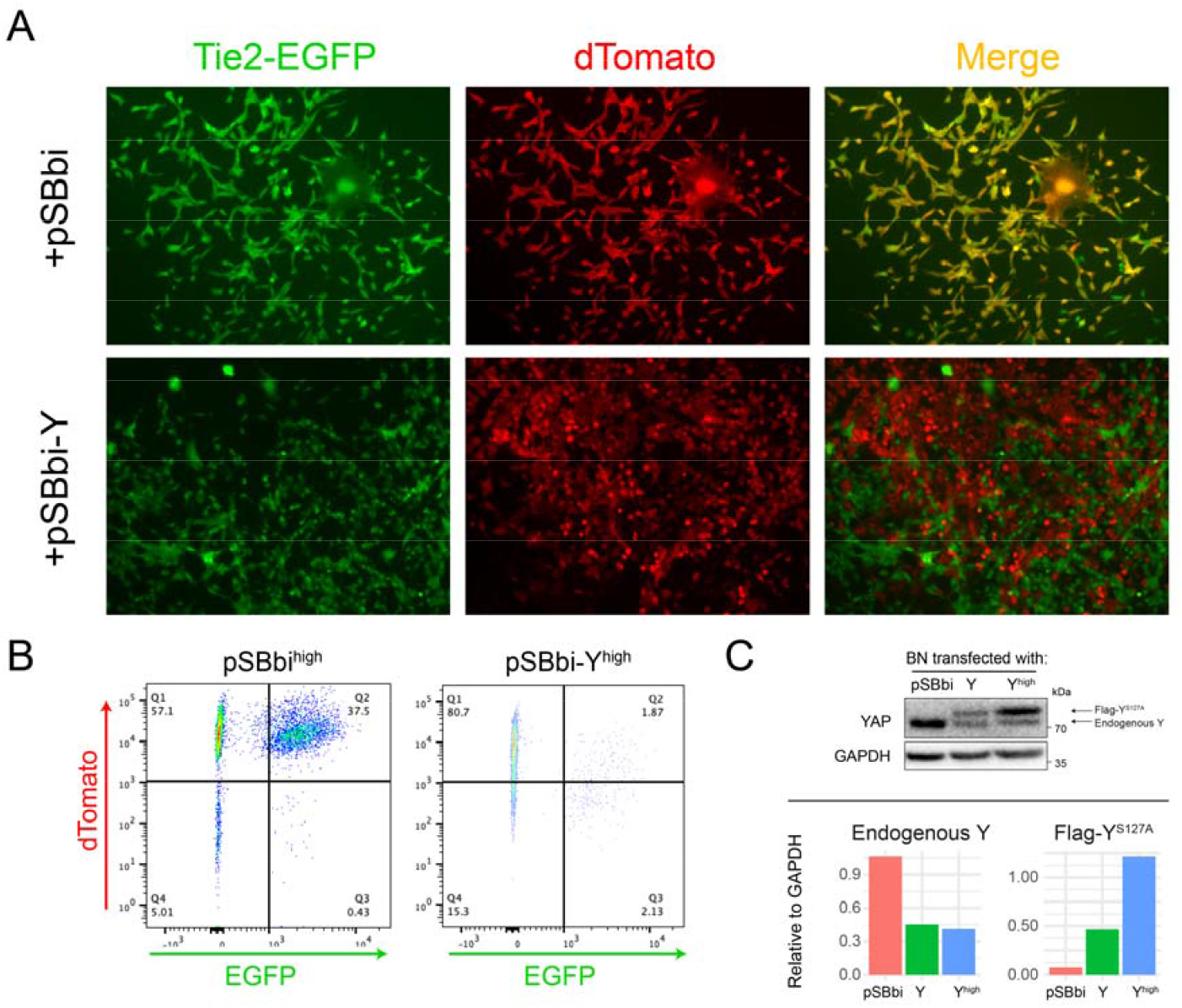
Exogenous Y over-expression suppresses Tie2-EGFP. **(A)** Following a 2-day period of hypoxia, BN1-Tie2-EGFP cells were sorted, expanded for 3-4 days and then transfected with a pSBbi-Y expression vector or the empty pSBbi vector as a control. Puromycin-resistant dTomato+ cells were then selected for 10 days and imaged by fluorescence microscopy. Merged images show that Y suppressed EGFP expression. **(B)** dTomato and EGFP expression in the cells from **(A)** were quantified by flow cytometry. **(C)** dTomato^high^ and unsorted BN1-Tie2-EGFP cells from **(B)** were expanded and examined by immunoblotting using an anti-Y antibody. Endogenous and exogenous Y were distinguished by virtue of the latter bearing a FLAG epitope and migrating more slowly.

### 3.7. HB cell lines have similar chemotherapeutic drug sensitivities

Current drug-based treatments for HB tend to be quite similar even though a number of studies suggest that certain molecular signatures may be predictive of chemotherapeutic responses and long-term survival [2,5,31,57-60]. This approach differs from that taken with many other human cancers whose underlying oncogenic drivers are often used to inform therapeutic options and stratify patients [61-66]. Having multiple immortalized and molecularly well-defined HB cell lines provided an opportunity to examine this question empirically. We therefore compared the sensitivities of representative BY, BYN, BN and YN cell lines to 4 drugs that are commonly used to treat HB, i.e. cis-platinum, etoposide, doxorubicin and vincristine. With few exceptions, all cell lines displayed similar sensitivities to each of these agents (Figure 8A and B). These studies thus indicate that the overall chemotherapeutic sensitivity of HB is little impacted by any of these 3 oncogenic mutations and in the face of cdkn2a locus inactivation.

**Figure 8.**
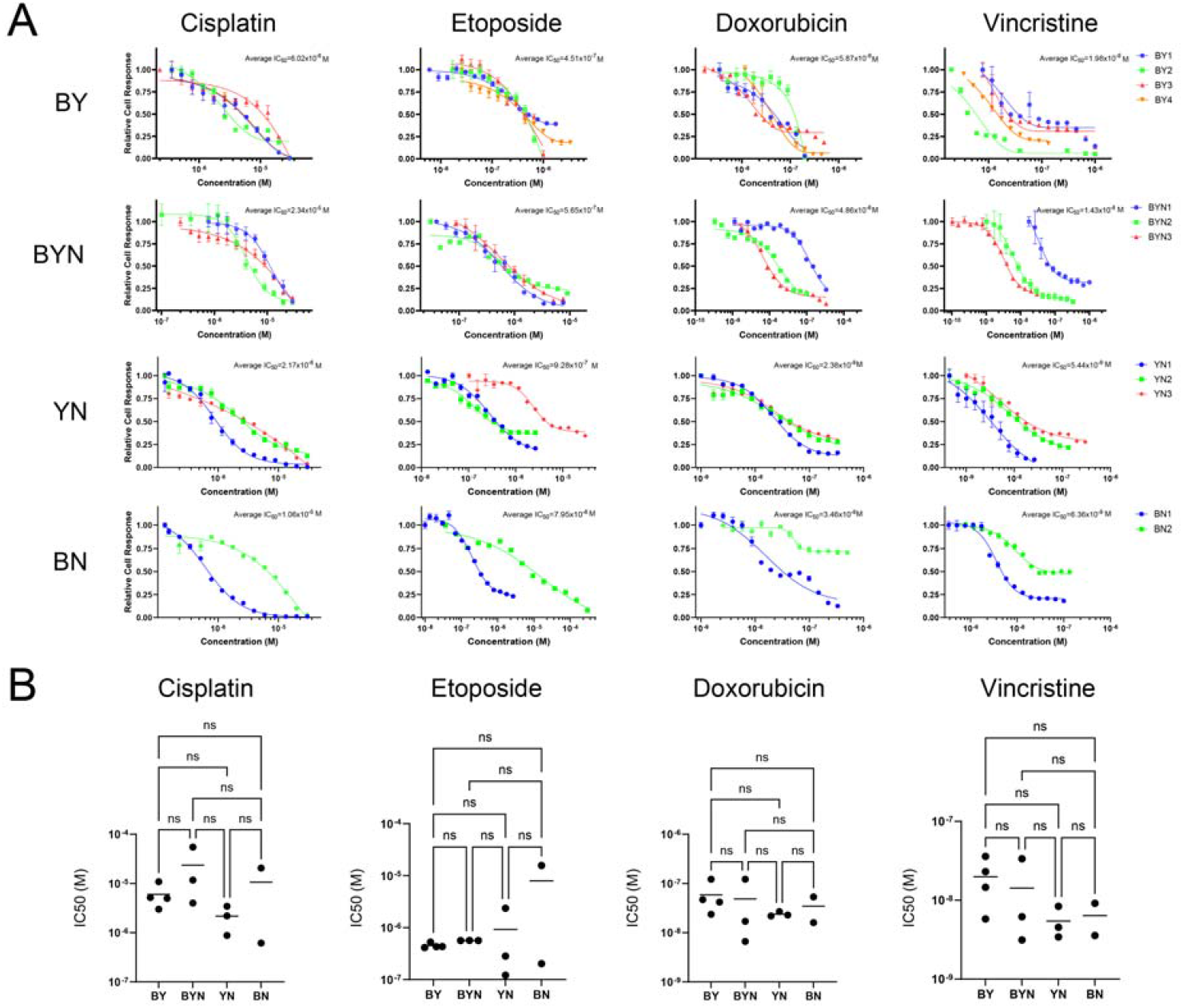
Murine HB cell lines display similar sensitivities to commonly employed therapeutic agents. **(A)** Typical dose-response curves. BY, BYN, BN and YN cell lines were seeded into 96 well plates, allowed to achieve log-phase growth over the next 24 hr and then exposed to the indicated concentrations of drugs for 72 hr. MTT assays were then performed, with each point represents the mean of 6 replicas +/-1 SE. **(B)** Chemotherapeutic drug sensitivities of all HB cell lines tested. Each point represents the calculated IC50 for one individual cell from the indicated molecular group based on dose-response profiles generated as described in **(A)**.

## 4. Discussion

Histologic, biochemical and molecular differences among HBs have allowed for various classification systems that are intended to identify tumors with favorable or unfavorable survival [2,5,9,20,22,45,46,57-60]. However, the tumor’s relative rarity, the fact that it seldom appears after the age of 4 years and that its overall survival/cure rate is ∼70% has made it challenging to conduct clinical studies aimed at individualizing chemotherapeutic regimens for these different groups while minimizing side effects, many of which are uniquely associated with this particularly young cohort [5,9,58,60,67,68].

The relatively small number of oncogenic drivers associated with HB classifies it as the least complex of human cancers [3,4,31,69,70]. Among the more common drivers, and associated with up to 80% of tumors, are mutations or amplification of *CTNNB1* and NRF2, unspecified defects in the Hippo pathway that increase Y’s nuclear accumulation and silencing of the CDKN2A tumor suppressor locus, typically by promoter methylation [6,9,15,25,59,70,71]. Indeed, any pairwise or triple combination of B, Y and N over-expression is sufficient to promote HB initiation in mouse models of the disease, with concurrent *Cdkn2a* mutations being needed to derive immortalized cell lines from the primary tumors as show both here and elsewhere [6,12,15,45,46]. However, this superficial assessment belies what is almost certainly a more complex and nuanced aspect of the disease. For example, it seems likely that different combinations and levels of expression of the above factors are responsible for different tumor histologies and behaviors, that different oncogenic B mutations drive distinct patterns of target gene expression and phenotypes and that much of HB pathogenesis and behavior remains reliant on less common driver mutations and/or incompletely understood epigenetic changes [9,11,12,25,31,57,58,69]. This combinatorial complexity, along with the acknowledged paucity of and pressing need for human HB cell lines [26,29] underscores the appeal of cell lines derived from molecularly defined murine tumors.

In the current work, we derived and characterized immortalized cell lines from primary BN and YN HBs [11]. As with BY and BYN cell lines, this required the Crisp/Cas9-mediated mutational targeting and partial inactivation of p16^INK4A^ and p19^ARF^ and their respective downstream retinoblastoma and TP53 TS pathways [24,26]. We have previously shown that all such mutationally-defined murine HBs and about half of human HBs somewhat surprisingly up-regulate WT p16^INK4A^ and p19^ARF^ (Figure 2B and ref. 26). We have interpreted this as evidence that, despite the high-level expression of these 2 potent TSs, it remains insufficient to override the potent proliferative stimuli originating from the various over-expressed oncogenic drivers. The mutant forms of p16^INK4A^ and p19^ARF^ that are expressed as a result of *in vivo* Crispr/Cas9-mediated targeting of the *Cdkn2a* locus (Figure 2A&B) likely represent less growth suppressive versions of their wild-type counterparts that were selected from amongst the large number of other mutant forms of these proteins that were generated (Supplementary File 1). While these various mutants remain capable of suppressing tumor cell growth, particularly when over-expressed (Figure 2E), they appear to be disabled enough to allow for immortalization and the establishment of permanent cell lines.

In addition to their tendency to recur, the most common p16^INK4A^ and p19^ARF^ mutations of BN and YN cell lines did not structurally resemble those of BY and BYN cell lines despite being generated in the same manner. This was supported by functional studies indicating that some mutants that suppressed BY and YN cell line proliferation did not suppress BN cells (Figure 2E). The differential selection of p16^INK4A^ and p19^ARF^ mutants among the cell lines likely reflects the degree to which they influence and/or are influenced by the combinatorial signaling of each of the B, Y and N TFs that are expressed by each of the 4 tumor groups [9,11].

A major rate-limiting determinant of a tumor’s growth, and in some cases an indicator of its aggressiveness, is the degree to which it establishes an independent vasculature to supply oxygen and nutrients and dispose of waste products [72,73]. The means by which this is achieved include the classical recruitment of pre-existing blood vessels from adjacent normal tissues (neo-angiogenesis), the formation of a *de facto* neo-vasculature directly from tumor cells themselves (vasculogenic mimicry) and the direct trans-differentiation of tumor cells into those resembling ECs [34,40-42,74-78]. With regard to the latter, we have previously demonstrated that BY cells can acquire EC-like characteristics but have shown here that this process is more impressive in BN cells (Figure 4). The acquisition of numerous EC-like properties is observed within 2 days following the exposure of BN monolayer cultures to hypoxia. Moreover, the subsequent formation of structures resembling EC-specific “tubes” appears to be quite robust as it occurs independent of the basement membrane substratum that is typically needed to support this state (Figure 4G) [79,80]. These attributes, which persist for several weeks following this single, transient hypoxic episode are accompanied by the expression of an array of EC-specific genes, many of which are expressed by both murine and human HBs (Figure 5A-D). In the former case, this is associated with expression patterns that clearly reflect the nature of the underlying molecular drivers (Figure 5A). This is in keeping with previous reports showing that EC-specific gene regulation is subject to control by the Wnt-β-catenin, Hippo and NRF2 pathways, whose terminal TFs not only include B, Y and N, respectively but also share considerable positive and negative cross-talk [11,81-83]. The greater variability of EC-specific gene expression by human HBs likely reflects greater overall molecular heterogeneity of these tumors as previously described (Figure 5C&D) [45,46,69].

It is of interest that in no cases do the EC-specific gene expression signatures of tu-mors or their cell lines resemble one another; nor do they precisely reflect those of normal livers, in which LSECs comprise 15-20% of the organ’s mass (Figure 5B-D and Figure 6G) [47,48]. However, the fact that the genewith their murine HB expression profiles of BN tumors and cell lines more closely resemble those of liver than do other HB cell types may explain why the former are much more predisposed to developing a more prominent vasculature (Figure 1B). The inability to do so in locations other than the liver may reflect the restricted and aberrant nature of their EC-specific gene expression that does not allow for trans-differentiation at other sites.

Importantly, the EC-like transcriptional profiles of our murine HBs and cells were not only distinct from one another but did not clearly recapitulate any particular known EC subtype (Figures 5 and 6G) [48,50-52].This, coupled with the fact that the above-mentioned Wnt-β-catenin, Hippo and NRF2 cross-talking pathways are aberrantly regulated in these cells, suggests that the observed EC-specific gene expression profiles represent abnormal hypoxic responses in which genes from multiple EC subclasses, are activated. As a result, only in some cases (e.g. in BN and less so in BY cells cells) [26] is the final collection of these transcripts sufficient and compatible to promote EC-like behaviors such as the ability to support a vasculature or tube formation (Figures 1A and 4G).

Differences in EC-specific gene expression were also noted in human cancers and the normal tissues from which they originated (Figure 6 I and J). As with their murine HB and HCC counterparts, none of the tumor-specific expression patterns resembled those of any normal tissues. Moreover, most tumor-specific expression patterns were also unique and could be further divided into subsets associated with significant survival differences (Figure 6K). This suggested that while many if not all cancers up-regulate subsets of EC-specific transcripts, only certain of these patterns confer a survival advantage. It remains to be determined whether these transcripts actually originate from tumor cells undergoing an actual transdifferentiation process such as we report here.

A potential explanation for why BN cells were particularly predisposed to EC-like trans-differentiation was suggested by the fact that they are the only one of our HB cell lines that does not express Y. Indeed, the enforced over-expression of Y in BN cells markedly inhibited Tie2-driven EGFP expression and down-regulated the expression of endogenous YAP (Figure 7C) [78,84]. In addition to this negative feedback control over its own promoter and that of its paralog, TAZ, YAP can either promote or inhibit EC differentiation in a context-dependent manner [13,53-56,74,78,82,84]. Interestingly, we did not find any evidence to indicate that human HBs with higher levels of EC-specific transcripts (Figure 5C and D) express less YAP or TAZ (not shown). Non-mutually exclusive reasons for this include the possibility that this regulatory pathway is not evolutionarily conserved, that human HBs are more molecularly heterogeneous or that EC-specific gene expression with actual functional consequences requires specific levels of β-catenin and/or NFE2 deregulation (i.e. different mutations or amplification versus mutation), which can exert significant effects on the overall transcriptional profile of the tumor [12,13,27,45,46,69,85]. The trans-differentiation of ECs from other tumor types such as gliomas and melanomas is well-established and may explain their resistance to anti-angiogenic agents that target proteins such as vascular endothelial growth factor, which recruit pre-existing exogenous ECs from sources residing in proximity to tumors [34,39,41,42,77,86,87].

A surprising finding from our studies was that BN1 cells exposed to hypoxia did not appear to retain their EC-like properties during *in vivo* tumorigenesis despite contributing extensively to growth and neo-vascularization (Figure 4H-J). Indeed, pure populations of EGFP+ BN1-Tie2-EGFP cells were able to generate HBs much more rapidly than control BN1 cells. This suggested not only that the EC phenotype was transient *in vivo* just as it is *in vitro* (Figure 4E and F) but that EC-like cells could revert back to tumor cells and generate tumors that were histologically indistinguishable from those formed by control BN1 cells (Figure 4I). This further suggested that the benefits conferred by EC-like BN1 cells likely occur early (i.e. prior to complete reversion back to tumor cells) perhaps by contributing to the early stages of tumor blood vessel formation and providing an environment that is more conducive to blood vessel formation by host-derived “true” ECs.

Although HB tends to respond well to chemotherapy and is associated with an overall long-term-survival/cure rate exceeding 70%, some tumors show poor responses to standard drug regimens or recur, in which case they are almost universally chemo-refractory [5,9,31,57,58,60,70]. Yet, despite certain pre-treatment molecular subtypes being associated with more unfavorable survival, current chemotherapeutic regimens do not account for these differences and thus do not stratify patients accordingly [7,57,59,60,69,70]. This is in marked contrast to many other cancers where certain molecular subtypes are often assigned to different treatment cohorts [65,66,88-90]. We took advantage of the well-defined and distinct molecularly properties of our 4 HB cell line groups [10,12,27] to ask whether they demonstrate any significant and consistent differences in their susceptibilities to chemotherapeutic drugs that are commonly employed to treat the disease. Although some differences were noted among individual cell lines within a specific molecular category, we were unable to make any significant inter-group distinctions (Figure 8). It is thus likely that factors other than B, Y and N are needed to impart the differential chemotherapeutic sensitivities that have been reported [5,57,58,69,70]. Important factors that are deserving of closer scrutiny include the degree to which the *CDKN2A* locus is inactivated given that the Rb and TP53 pathways that are regulated by p16^INK4A^ and p19^ARF^ are major determinants of both survival and chemotherapy responses [23-25]. It remains possible that are cell lines will nonetheless be useful in testing of new chemotherapeutic agents as they emerge.

## 5. Conclusions

The work presented here and elsewhere [26] provides a means by which murine HB cell lines, with defined molecular drivers akin to those of their human counterparts, can be reproducibly isolated from primary tumors. In the vast majority of cases, they remain tumorigenic in immune-competent hosts, are capable of forming metastasis-like pulmonary lesions and retain the histologies of the tumors from which they originate. As such, they provide a much-needed set of well-defined reagents that have heretofore been lacking in the study of this disease [6,29]. We envision these cell lines as having future utility in the study of tumor metabolism, the trans-differentiation of tumor cells to ECs and drug screening. The ability of BN cells to acquire EC-like properties in response to hypoxia also provides a new model in which to more closely study the trans-differentiation process and the molecular events that are responsible for this reversible transition.

## Supporting information

Supplementary File 3. List of the unique subset of EC-specific genes expressed in normal murine livers (Figure 5B). These are likely expressed by live

Supplementary file 4. List of the 200 hypoxia-responsive genes shown Figure 6C.

Supplementary file 5. List of the 194 liver-specific genes shown in Figure 6E.

Supplementary File 1. The most commonly identified Cdkn2a exon 2 mutations and their frequencies in immortalized BY and BN cell lines. Structures of t

Supplementary File 2. List of the 1853 EC-specific, non-redundant murine genes from Figure 2B.

## Supplementary Material

The following supporting information can be downloaded at: https://www.mdpi.com/article/doi/s1, Figure S1: Additional examples of H&E-stained sections of subcutaneous tumors generated by the the indicated cell lines; Figure S2: Additional examples of H&E-stained sections of tumors generated in lungs by the indicated cell lines following tail vein injection. Table S1: The most commonly identified *Cdkn2a* exon 2 mutations and their frequencies in immortalized BY and BN cell lines. Table S2: list of the 1853 EC-specific, non-redundant murine genes from Figure 2B. Table S3: List of the unique subset of EC-specific genes expressed in normal murine livers. Table S4: List of the 200 hypoxia-responsive genes shown Figure 6C. Table S5: List of the 194 liver-specific genes shown in Figure 6E.

## Author Contributions

Conceptualization, EVP and HW; methodology, CK, CH, JK HW, EVP; validation, KC, AT, HW, EVP; formal analysis, KC, SR, HB, EVP; investigation, KC, AT, CH, JK, JL, SR, HW; resources, EVP; data curation, KC, HW.; writing—original draft preparation, EVP; writing—review and editing, EVP, KC, HW.; visualization, KC, HW.; supervision, HW, EVP.; project administration, EVP; funding acquisition, EVP. All authors have read and agreed to the published version of the manuscript.

## Funding

Funding for this research was in part provided by the UPMC Children’s Hospital of Pittsburgh Research Advisory Committee, The UPMC Children’s Hospital of Pittsburgh Foundation and The Rally Foundation. This research was supported in part by the University of Pittsburgh Center for Research Computing through the resources provided. Specifically, this work used the HTC cluster, which is supported by NIH award number S10OD028483

## Institutional Review Board Statement

The animal study protocol was approved by the Institutional Review Board (or Ethics Committee) of The University of Pittsburgh (protocol code 23083650, approved 8/16/2023).

## Data Availability Statement

RNA-seq data have been deposited in the NCBI Gene Expression Omnibus (GEO) under accession number GSE302803 and can be accessed at: https://www.ncbi.nlm.nih.gov/geo/query/acc.cgi?acc=GSE302803. Some of the result reported here relied upon data generated by the TCGA Research Network: https://www.cancer.gov/tcga (ac-cessed on May 4, 2025). All other data supporting the findings of this study are available from the corresponding author upon reasonable request..

## Acknowledgments

We thank Dr. Jianhua Luo and Dr. Silvia Liu of the Pittsburgh Liver Research Center’s Genomics and Systems Biology Core for their assistance with RNA-seq.

## Conflicts of Interest

The authors declare no conflicts of interest.

## Abbreviations

B: a mutant of human β-catenin (Δ90)
BN: HBs induced by over-expression of B and N
BY: HBs induced by over-expression of B and Y
BYN: HBs induced by over-expression of B, Y and N
EC: endothelial cell
EGFP: enhanced green fluorescent protein
GSEA: gene set enrichment analysis
HB: hepatoblastoma
HCC: hepatocellular carcinoma
HDTVI: hydrodynamic tail vein injection
LESC: liver sinusoidal endothelial cells
N: a mutant of YAP (Y^S127A^)
NY: HBs induced by over-expression of N and Y
SB: Sleeping Beauty plasmid vector
TF: transcription factor
TS: tumor suppressor
WT: wild-type
Y: a mutant of NRF2 (N^L30P^)

## Notes

### Competing Interest Statement

The authors have declared no competing interest.

