## Supplementary file legend for "Derivation of Genetically-Defined Murine Hepatoblastoma Cell Lines with Angiogenic Potential"

**Supplementary Files**

**1. Supplementary File 1. The most commonly identified *Cdkn2a* exon 2 mutations and their frequencies in immortalized BY and BN cell lines.** Structures of the top 5 from each cell shown are shown in Figure 2A.

**2. Supplementary File 2.** List of the 1853 EC-specific, non-redundant murine genes from Figure 2B.

**3. Supplementary File 3.** List of the unique subset of EC-specific genes expressed in normal murine livers (Figure 5B). These are likely expressed by liver sinuosoidal ECs (LSECs) that normally comprise 15-20% of the organ’s cellular mass (42,43).

**4. Supplementary file 4.** List of the 200 hypoxia-responsive genes shown **Figure 6C.**

**5. Supplementary file 5.** List of the 194 liver-specific genes shown in **Figure 6E.**
